# Plants cultivated for ecosystem restoration can evolve towards a domestication syndrome

**DOI:** 10.1101/2023.02.27.530164

**Authors:** Malte Conrady, Christian Lampei, Oliver Bossdorf, Norbert Hölzel, Stefan Michalski, Walter Durka, Anna Bucharova

## Abstract

The UN Decade on Ecosystem Restoration calls for upscaling restoration efforts, but many terrestrial restoration projects are constrained by seed availability. To overcome these constraints, wild plants are increasingly propagated on farms to produce seeds for restoration projects. During on-farm propagation, the plants face non-natural conditions with different selection pressures, and they might evolve adaptations to cultivation that parallel those of agricultural crops, which could be detrimental to restoration efforts. To test this, we compared traits of 19 species grown from wild-collected seeds to those from their farm-propagated offspring of up to four cultivation generations, produced by two European seed growers, in a common garden experiment. We found that some plants rapidly evolved across cultivated generations towards increased size and reproduction, lower within-species variability, and more synchronized flowering. In one species, we found evolution towards less seed shattering. These trait changes are typical signs of the crop domestication syndrome, and our study demonstrates that it can also occur during cultivation of wild plants, within only few cultivated generations. However, there was large variability between cultivation lineages, and the observed effect sizes were generally rather moderate, which suggests that the detected evolutionary changes are unlikely to compromise farm-propagated seeds for ecosystem restoration. To mitigate the potential negative effects of unintended selection, we recommend to limit the maximum number of generations the plants can be cultivated without replenishing the seed stock from new wild collections.

**Significance Statement:** Globally upscaling demands for native seeds for ecosystem restoration can be covered by agricultural seed propagation. Yet, agricultural practice can unintentionally select for specific traits and reduce adaptive variability, which could affect plant performance once sown back to the wild. We show, across 19 wild species, two seed producers and up to four consecutive cultivated generations, that some plants under cultivation evolved higher vigor, reduced adaptive variability, synchronized flowering and in one case, reduced seed shattering. Yet, there were substantial differences among cultivation lineages, with negligible changes in most, and large changes only in a few cases. Substantial unintended evolution in cultivation is thus rather an exception than the rule.

## Introduction

Ecosystem restoration is increasingly recognized as an indispensable tool to address the current biodiversity and environmental crisis (1). However, many degraded terrestrial ecosystems lack sufficient diaspores for the regeneration of native vegetation, and therefore restoration often relies on the introduction of plants from other sources (2). While forests are usually restored by planting nursery-grown seedlings, grasslands and drylands are restored by direct seeding – an approach that requires large seeding densities since the establishment success of sown seeds is often low (3). This causes an unprecedented demand for native seeds, particularly in the context of the global restoration movement (www.decadeonrestoration.org), making seed availability a major bottleneck for upscaling restoration (4, 5). Wild-collected seeds cannot cover the demand because large-scale seed harvesting from natural populations threatens the persistence of the donor populations (6, 7). Consequently, wild-collected seeds are increasingly used for large-scale seed propagation on farms where plants are grown as crops, and their seeds are then used for restoration projects (8, 9).

Plants that are grown in agricultural propagation face novel, unintended selection pressures. The propagation of seeds for ecological restoration generally aims to maintain the natural properties of wild populations, such as large phenotypic and genetic variation (10, 11). This is different in agricultural cultivars, which are intentionally bred for specific traits like large size or high seed production but with low phenotypic diversity (12). Even when there is no such intentional selection, cultivation processes might impose unconscious selection, shifting plant traits towards cultivation-specific adaptation, and reducing within-population phenotypic variability (10). Similar to early crop domestication, where humans unconsciously selected for certain phenotypes (13, 14), this may result in a set of traits referred to as domestication syndrome. This includes taller growth, increased apical dominance, synchronized flowering, higher seed investment, reduced seed shattering, and a number of physiological traits, including the loss of dormancy (15–17). As a result, domesticated crops are well adapted to cultivation but can only rarely survive in the wild (15). If wild plants cultivated for ecological restoration underwent similar domestication processes, the resulting seeds could be poorly adapted to natural conditions and thus unsuitable for restoration.

The potential evolution of wild plants during cultivation for ecological restoration has been intensely debated, but experimental evidence is scarce and inconclusive (10, 11). Some previous studies used molecular markers to understand genetic drift during the seed production (18, 19), yet none of these focused on adaptive genetic variation and thus, were unable to test for selection (20). Common garden studies that tested for heritable phenotypic changes were so far limited to individual species, only one cultivated generation, or they lacked adequate comparisons to the wild ancestors (21–23), and the obtained results vary. To assess how common and how severe evolution is during seed propagation for ecosystem restoration, we need systematic information on many species, ideally from multiple seed production systems.

Here, we focused on evolution during seed production in 19 perennial species. All species are native to European mesic grasslands, and although some of them behave as invasive weeds in other parts of the world, they are not weedy in their native habitats. We received the seed material from two European seed producers, one located in Germany and one in Austria (hereafter called Producer 1 and Producer 2, Table 1, Figure 1). Because the seed producers archived a sample of each seed lot, we were able to obtain wild-collected seeds (F0) that were used for the establishment of each cultivation lineage, as well as up to four consecutive generations that were produced from these wild seeds (F1-F4, Table1). Seven species were available from both seed producers and one species from two regions from one seed producer, resulting in a total of 27 independent cultivation lineages (i.e. F0 plus the corresponding cultivated generations) and a total of 93 generations. To test for heritable adaptive differentiation across generations in cultivation, we grew plants from several consecutive generations of farm-grown seeds (F1-F4) side by side with their wild-collected ancestors (F0) in a common garden and recorded plant height, the number of flowers, aboveground biomass, phenology and, in species where the morphology allowed this, seed shattering. We focused on adult traits because earlier plant life-stages are more likely to be affected by maternal effects (24). We hypothesized that across cultivated generations, (1) plants evolved towards taller growth and higher biomass, higher seed investment, earlier flowering and lower seed shattering, (2) the variation in traits and phenology generally decreased and (3) that the magnitudes of evolutionary changes depended on species and/or cultivation lineage.

**Table 1.** Species and their cultivated generations included in the common garden experiment. F0 are the wild collected seeds, and F1-F4 are consecutive generations in cultivation.

| Species | Family | Producer 1 | Producer 2 |
| --- | --- | --- | --- |
| *Achillea millefolium* | Asteraceae | F0-F2 | F0; F3; F4 |
| *Anthoxanthum odoratum* | Poaceae |  | F0-F2 |
| *Centaurea jacea* | Asteraceae | F0-F3 |  |
| *Centaurea scabiosa* | Asteraceae | F0-F2 | F0-F3 |
| *Crepis biennis* | Asteraceae | F0-F3 | F1-F3 |
| *Cynosurus cristatus* | Poaceae |  | F0-F2 |
| *Dianthus carthusianorum* | Caryophyllaceae |  | F0-F4 |
| *Galium verum* | Rubiaceae |  | F0-F2 |
| *Leontodon hispidus* | Asteraceae | F0-F2 | F1-F3 |
| *Leucanthemum ircutianum* | Asteraceae | F0-F3 | F0-F2; F4 |
| *Lotus corniculatus* | Fabaceae | F0-F3 | F0; F1 |
| *Lychnis flos-cuculi* | Caryophyllaceae |  | F0-F2 |
| *Ranunculus acris* | Ranunculaceae | F0-F3 |  |
| *Rumex acetosa region R6* | Polygonaceae | F0-F2 |  |
| *Rumex acetosa region R16* | Polygonaceae | F0-F2 |  |
| *Salvia pratensis* | Lamiaceae |  | F0; F1; F3 |
| *Silene dioica* | Caryophyllaceae | F0-F3 |  |
| *Silene vulgaris* | Caryophyllaceae | F0-F3 | F1-F4 |
| *Trifolium pratense* | Fabaceae | F0-F3 |  |
| *Veronica teucrium* | Plantaginaceae |  | F0-F2 |

**Figure 1.**
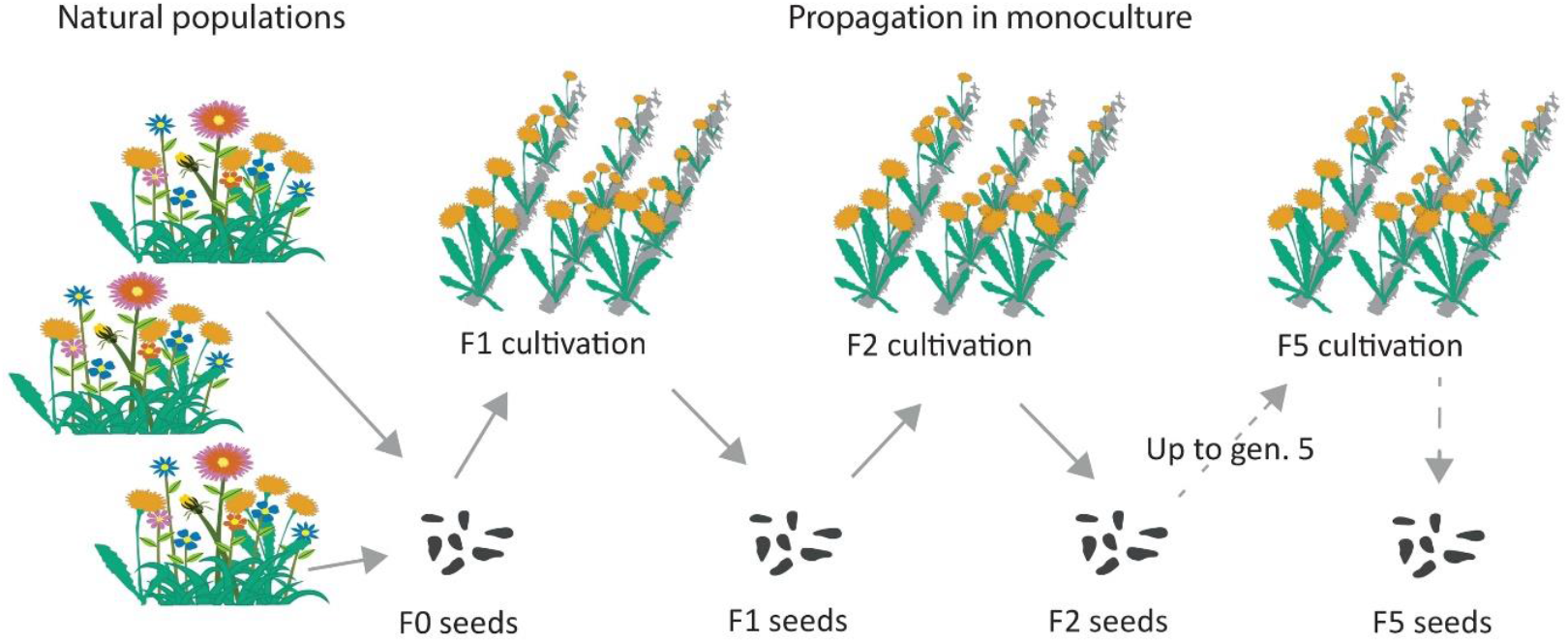
Process of propagation of wild plants for ecosystem restoration. The F0 seeds are collected from several (Producer 1) or one (Producer 2) large natural population. On farm, seeds are propagated in monoculture and the farm-produced seeds are available for restoration projects, with a small part being used to establish the next cultivated generation. The process can be repeated for up to five cultivated generations; then the seed stock must be replenished from new wild collection.

## Results

First, we tested for general patterns of evolution under cultivation across all species, cultivation lineages and generations, relating trait values to the generation number (F0-F4) in linear mixed models. We also estimated variation within generations of each cultivation lineage, expressed as coefficient of variation, and related it to the generation number. We found that across generations in cultivation, aboveground biomass increased, plants produced more flowers and were increasingly taller. There was no significant trend in the start of flowering (Figure 2, Table S2). The variation in trait values decreased across cultivated generations for aboveground biomass, the number of flowers and especially for the start of flowering, suggesting loss of functional genetic variation and evolution towards synchronized flowering (Figure 2, Table S2).

**Figure 2.**
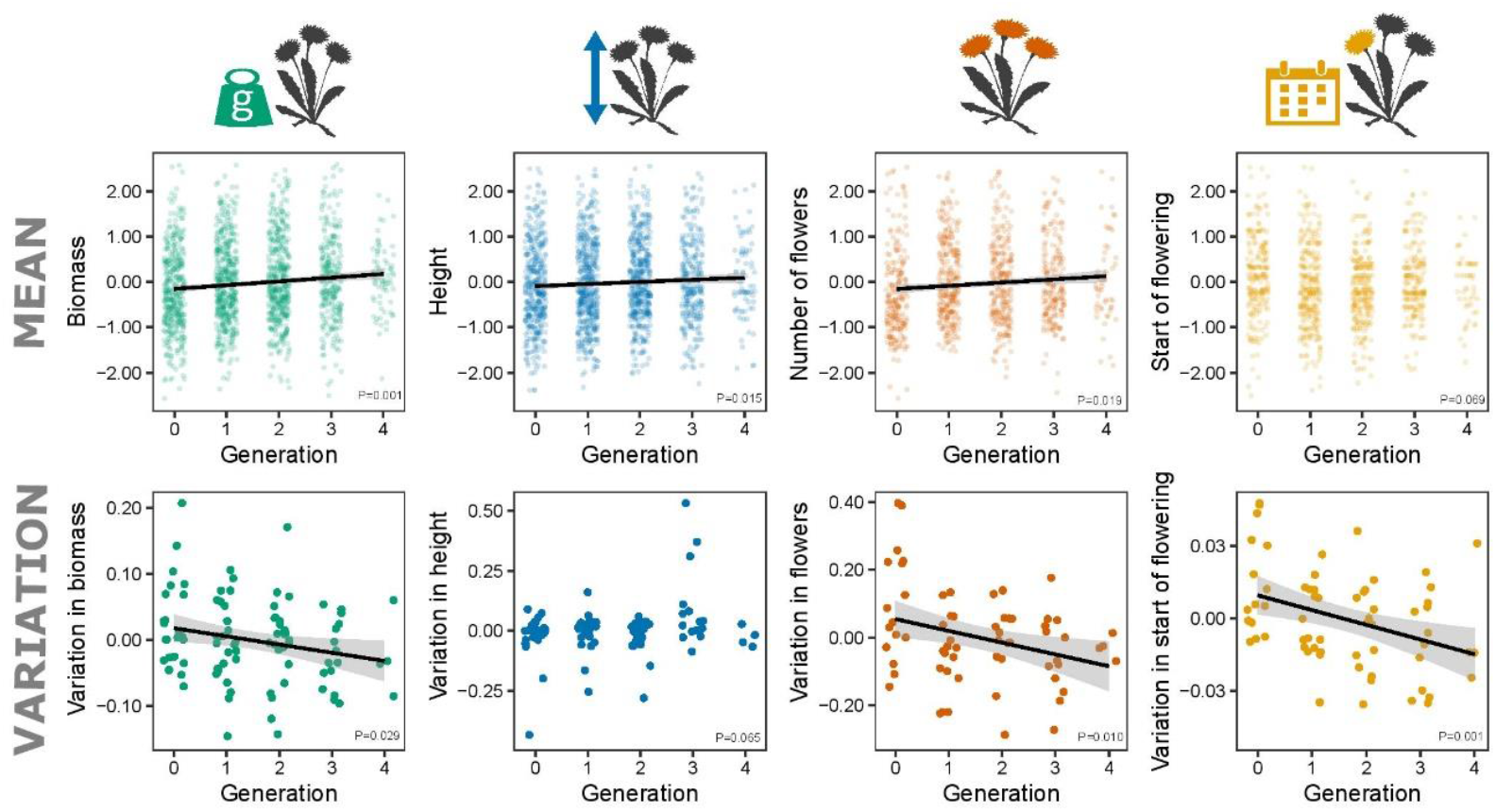
Analyses across all cultivation lineages. Top row: Changes of biomass, height, number of flowers and the start of flowering across generations in cultivation. Each point is an individual plant, with values scaled within cultivation lineages. Bottom row: the corresponding changes of within-generation coefficient of variation across generations in cultivation. Each point is the coefficient of variation of one generation and cultivation lineage; the values are adjusted for cultivation linages. Lines indicate significant relationships. See Table S1 for model results.

Second, to understand how evolution under cultivation varies among species and cultivation lineages, we analyzed the data for each cultivation lineage separately. Although the majority of changes within cultivation lineages pointed in the same direction as the overall trends, i.e. plants were getting larger, taller and had more flowers across cultivated generations, less than half of the individual cultivation lineages showed a significant change in at least one trait (11 out of 27, Figure 3). On average, the trait changes per generation were 6.7% of the initial trait range detected in plants from wild-collected seeds (Table S2). Nevertheless, in several cultivation lineages, we detected substantial changes. For example, in *Lychnis flos-cuculi*, aboveground vegetation increased by 24%, number of flowers by 34% and the start of flowering shifted seven days earlier per generation. In *Leontodon hispidus* (the lineage from Producer 1), biomass and number of flowers increased by 17% and 43% per generation, respectively. However, such strong effects were rare (Figure 1, Table S2).

**Figure 3.**
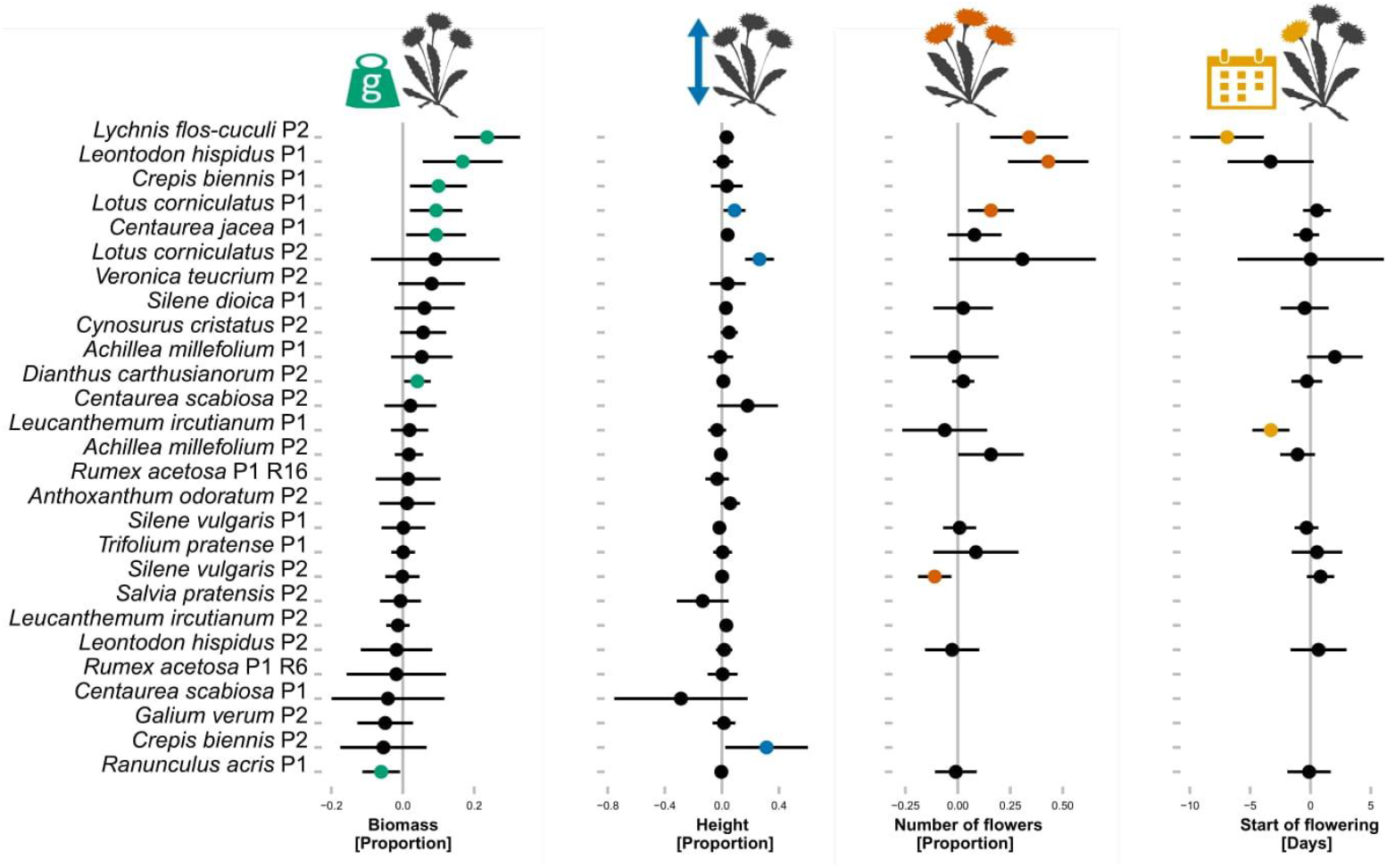
The changes in biomass, height, number of flowers and start of flowering across cultivated generations in 27 cultivation lineages. The values are effect sizes and their confidence intervals. The effect sizes for number of flowers and start of flowering were calculated only for cultivation lineages where 50% of individuals started to flower during the experiment. The effect sizes for biomass, height and number of flowers are expressed as proportion of the mean value of a given cultivation lineage, with positive values indicating increase across generations. The effect sizes for start of flowering are in days, with positive values indicating later flowering. Colored symbols are effects significant at *P* < 0.05. See Table S2 for model results.

Third, we focused on the reduction of seed shattering as a prominent trait of the domestication syndrome. This was possible in only three species (*Centaurea jacea, Lotus corniculatus* and *Silene vulgaris*) in which there was variability in this trait at the time of the harvest, and the inflorescence morphology allowed reliable estimation. For each plant, we recorded how many ripe flowers still had the seeds attached and how many had lost the seeds, and expressed it as odds ratio. While we detected no significant change across generations in *Lotus corniculatus* and *Silene vulgaris*, the proportion of flowers with attached seeds increased across generations in *Centaurea jacea* (odds ratio 1.36 per generation, Figure 4, Table S2).

**Figure 4.**
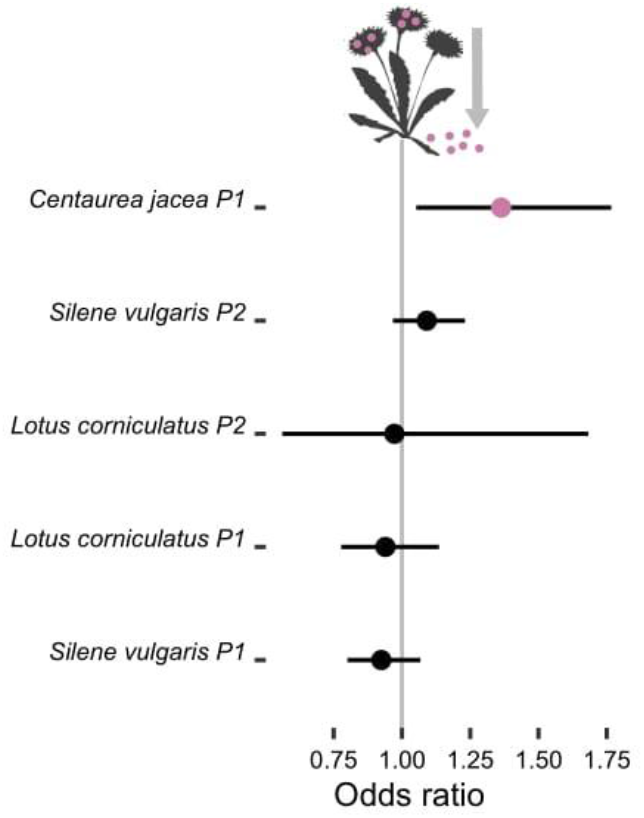
Changes in retention of seeds across generations, the values are effect sizes and their confidence interval. The colored point indicates an effect significant at *P* < 0.05. Effect sizes (with a confidence interval) are odds ratios for flowers with seeds versus flowers without seeds, with values >1 indicating increased seed retention. See Table S2 for model results.

## Discussion

The propagation of native plants for seed production is indispensable to ensure sufficient seeds for upscaling restoration efforts and counteracting biodiversity declines (11). Yet, agricultural cultivation bears a risk of unintended selection, which could alter adaptive traits and reduce plant adaptation to natural conditions (10). Here we show, across 19 species, multiple cultivation lineages and two European seed producers, that plants from farm-propagated seeds evolved towards a domestication syndrome across as few as 2-4 cultivated generations. However, the rate of change varied between species and cultivation lineages, with substantial changes in only a handful of lineages while the majority showed little or no change.

Plants grown from farm-propagated seeds were increasingly larger, taller, produced more flowers, and became less variable in these traits the more generations they spent in cultivation. These increases in size and vigor may appear beneficial at first, as more seeds are available for restoration. However, plant vigor can be selected against in natural populations because of trade-offs between vigor and drought tolerance or herbivory resistance (25–27). Agricultural cultivation often involves watering and protection from herbivores, which relaxes selection for drought and herbivore resistance, and it may favor high-vigor plants, which produce more seeds and therefore contribute more to the next generation. The increase of vigor traits was accompanied by a reduction of their variability, which could be attributed to less effective seed harvesting from genotypes that deviate too much from the mean. Both increase in vigor and loss of variation are typical elements of the domestication syndrome, i.e. when early crops adapted to agricultural conditions (15).

Across generations in cultivation, wild plants evolved towards more synchronized flowering phenology and, in one case, retention of ripe seeds – also well-known symptoms of the cultivation syndrome (13). In wild populations, the fitness of a particular genotype is determined by the number of ripe seeds that fall on the ground and eventually germinate in the next generation. In agricultural propagation, only the seeds that are ripe at the time of harvest can be harvested and thus contribute to the next generation, which excludes both early- and late-flowering genotypes and synchronizes the flowering time of propagated plants. Additionally, crops evolved retention of ripe seeds, in contrast to wild plants which typically disperse seeds as soon as they are ripe (28). We detected this phenomenon in *Centaurea jacea* where later generations had more flowers with non-dispersed seeds at the time of harvest, even though the start of flowering did not shift across cultivated generations in this species.

The observed evolution of some wild plants towards a domestication syndrome was rapid, within only few cultivated generations. So far, the domestication syndrome was studied on the scale of hundreds to thousands of years, by comparing crops with their wild relatives, or testing for evolutionary signals in genomes of current or ancient crops (e.g. (15, 28–30). Our study system allowed us to reconstruct evolution in response to cultivation, from generation to generation. It thus provides unique data on how rapidly this evolution can act. Similar to ancient domestication (13, 14), there was no intentional selection in our system, as the seed producing farmers do not breed for specific traits and in fact try to avoid selection (31). The observed changes thus can be attributed to unintentional selection exerted by agricultural practices. The high rate of evolution of some species is not surprising. Rapid evolutionary responses to novel environments across few generations was shown before in response to climate change, seeding to a degraded habitat or exposition to a novel pollinator community (32–34).

The direction of evolution in individual cultivation lineages mostly pointed in the same direction as the general trend, yet the rates of change varied. While more than half of the lineages did not significantly change in any of the studied traits, some lineages changed strongly. The rate of change was independent of species identity: for example, *Leontodon hispidus* from Producer 1 underwent substantial changes in biomass and number of flowers, while the same species from Producer 2 did not change in any of these traits. This suggests that the rate of change cannot be predicted from species identity or plant traits. Instead, the magnitudes of changes likely depend on seed production practices and individual decisions during the production process. For example, the largest rate of change was detected in *Lychnis flos-cuculi*, both in vigor and flower phenology. The farmer producing this specific cultivation line harvests the seeds when they subjectively estimate that the majority of plants bear ripe seeds (*pers. comm*.). Yet, this approach effectively selects against late flowering genotypes and shifts the population mean towards early flowering. The earlier flowing genotypes are also more vigorous in this species (Figure S1), which contributes to the shift in the vigor-related traits. Such details of production steps, specific to individual farms and cultivation lineages, are probably driving many of the observed differences in rates of change.

The results of our study may, to some degree, depend on the life-histories of our species. We studied perennial species of mesic grasslands with rather short generation cycles. In other habitats and species with different life histories, the strength of selection and impact of cultivation may differ. However, longer generation cycles do not necessarily imply different rates of changes, because we described changes per generation, not per years. However, if longer-lived species harbor more within-population variability in age of first reproduction, there will be particularly strong selection for early reproducing genotypes, because farmers will start to harvest as soon as the first genotypes produce seeds. Moreover, in plants with strong dormancy, as is common in drylands (35), there might be additionally strong selection for loss of dormancy. Espeland et al. (10) elegantly summarized how plant life histories can affect the strength of evolution during cultivation, yet experimental tests are missing so far. To address these questions, future research should include species with a broader spectrum of life histories and habitats of origin.

Our study covers many species and presents data on evolutionary changes from generation to generation, filling an important gap in previous research. However, this strength necessarily comes with some limitations, because there is almost always a trade-off between generality and precision (36). For instance, the seeds we used differed in age and possibly in maternal environment, which could affect plant phenotypes through maternal effects. However, while maternal effects may be present, we have reasons to doubt that they are the cause for the detected trends. First, we constrained our analyses to adult traits which are generally less likely to be affected by maternal effects than early traits such as seed dormancy, germination or early growth (24). Second, maternal effects have been found to be weaker in perennial plants than in annuals (37). Third, maternal effects tend to particularly appear under stress, but are often less visible under favorable conditions (38). Fourth, the direction and strength of maternal effects varies among species and even genotypes of the same species (39, 40) and therefore may simply add noise to our observations from genetically diverse cultivation lineages of many species (18). Finally, it is not very likely that maternal effects from different years would induce a linear trait change across generations. Thus, selection through continuous cultivation appears a much more parsimonious explanation.

### Implication for practice

Agricultural propagation of native plants is mandatory to provide a sufficient amount of seeds for upscaling ecological restoration (11). However, there are concerns that plants could adapt to cultivation and lose adaptation to natural conditions, which could be detrimental for the restoration success (10). We indeed detected substantial changes in several cultivation lineages, for example one week earlier start of flowering per generation or 17% increase in number of flowers per generation. These traits are commonly adaptive in the wild, and it is thus likely that cultivation affects performance of the plants in restoration sites. Nevertheless, such large changes were exceptional. In the majority of cultivation lineages, we did not detect any significant change in any of the measured traits, and the average rate of change per generation was some 7% of the trait value range present in the wild populations. However, some evolutionary changes may only become visible when plants face stress (e.g. drought, herbivory), and the benign conditions in our study may have underestimated the evolutionary potential to some extent. Some previous studies (22, 23) reported massive evolutionary change in response to cultivation of individual species, yet such effects seem to be an exception rather than a rule, at least in the system we studied.

We detected a slight loss of adaptive heritable variability across cultivated generations. This is worrisome because reduced variability could limit the ability of restored populations to adapt to their novel habitats, or other environmental changes (10). However, the reduction of variability was rather moderate, and appears unlikely to have detrimental effects on the adaptive potential – at least across the first few cultivated generations.

The extent of unintended evolution during cultivation may be mitigated by adjustments of production methods and procedures. Reduction or temporary cessation of watering and herbivore control could reduce the advantage of vigorous plants and thus reduce selection for vigor. Harvesting ripe seeds multiple times per season could reduce shifts in flowering time and synchronization. However, such measures require increased effort, and they may reduce seed yield. As seed availability is one of the main limiting factors for restoration globally (4, 5), upscaling seed production has become an important goal (11). Implementing measures that complicate the seed production process and potentially reduce seed yield might be counterproductive in this context. Measures are needed particularly in those few cultivation lineages with documented large trait shifts, but in the majority of cases the absolute changes were very moderate across the first few generations and, in our opinion, tolerable. After a few generations in cultivation, agriculturally produced seeds of wild plants are still regionally adapted (41). As trait variability was only moderately reduced by cultivation, plants from cultivated seed should still harbor substantial genetic diversity and thus be able to adapt to the novel conditions at the restoration site. This may eventually override the minor changes caused by cultivation (34). However, even a minor incremental increase could yield substantial change if a seed stock was cultivated for many generations. We thus strongly recommend to limit the maximum number of generations a seed stock can be cultivated without replenishing from new wild collection, and follow the example of *Regiosaatgut* in Germany (5 generations allowed, (31)) or Yellow Tag in the US (3-5 generations (42)).

## Materials and Methods

### Seeds

We used seeds from two seed producers – further called Producer 1 and Producer 2, one located in Germany and one in Austria. The producers are larger companies that have contracts with many small farmers who produce the seeds on their farms. The production starts with seed collection from multiple (Producer 1) or one (Producer 2) large natural populations (Figure 1) (31). This wild-collected seeds (F0) are then propagated in a horticultural setting and the plugs are transferred to fields. Seeds of this first cultivated generation (F1) are partly used for restoration projects and partly for establishing the next generation in cultivation (F2). The process can be repeated for up to five cultivated generations (F5), then, to prevent genetic deterioration, the seed stock must be replenished with new wild collections. We obtained seeds of 27 independent cultivation lineages from 19 species, which consisted of archived wild collections and up to 4 consecutive cultivated generations (for six species we had two cultivation lineages). In total we had 93 accessions (individual generations of each cultivation lineage) (Table 1).

### Common garden experiment

In March 2018, we planted 33 seeds per accession into seeding trays, with 3 replicates, and placed them in a cold greenhouse. When the seedlings developed their first true leaves, we transplanted 26 seedlings per accession individually to quickpots, and after three further weeks we planted 18 of these (randomly selected) into 2.5 l pots filled with standard potting soil. We placed the pots in a common garden, in a randomized blinded design, and watered them as needed. During the entire experiment, we recorded the start of flowering three times a week. After three months, we measured plant height and counted the number of inflorescences (for the sake of simplicity called flowers in this paper). We harvested all aboveground biomass, dried it for 48 hours at 70°C, and weighted it. In three species (*Centaurea jacea, Silene vulgaris* and *Lotus corniculatus*), we estimated the odds ratio of seed shattering as the number of flowers with ripe seeds still attached to the plant versus the number of flowers that dropped the seeds.

The seeds we used in this experiment differed in age and possibly in maternal environment, which could both affect plant phenotypes. However, standardizing the seed quality through growing an intermediate refresher generation (e.g. (32, 43)) was not possible due to the high number of species and populations, and their outcrossing and perenniality (not all species flower in the first year). To avoid strong influences of maternal effects we constrained our study to traits of adult plants.

### Data analysis

All analyses were performed in R (R Development Core Team, 2021). First, we focused on the effect of cultivation on start of flowering, total biomass, height and number of flowers across all cultivation lineages. We related each of the four response variables to generation as a continuous explanatory variable and the cultivation linage as random factor in a mixed model. The term cultivation lineage was approximately equivalent to species nested within seed producer. However, in one species we had two cultivation lineages each from two regions from the same seed producer (Table 1). Including producer, species and region as random factors resulted in such a complex model structure that the model did not converge. We thus included only cultivation lineage as a random factor. A model with species nested within producer as random factors yielded nearly identical results. To allow cross-lineage comparison, all response variables were scaled within cultivation lineage. Additionally, we tested whether the variation in these traits changed across generations. We calculated the coefficient of variation for each plant trait in each generation in each cultivation lineage, excluding accessions where traits were available from less than four individual plants. We then related the coefficients of variation in each trait to the generation as continuous explanatory variable and cultivation line as a random factor. In both analyses, we used the R-package *nlme* (44). We did not analyze seed-shattering in a cross-lineage analysis because this trait was available only for five cultivation lineages of three species.

Second, we tested for changes in biomass, height, number of flowers, start of flowering and seed shattering across generations in individual cultivation lineages. For each trait and cultivation line we fitted a linear model that had one of the above-mentioned traits as a response variable and the generation number as continuous explanatory variable. For biomass and height, we ran these models for all 27 cultivation lineages. For the number of flowers and start of flowering, we analyzed only the 15 cultivation lineages where at least 50% of individuals had started flowering. Seed shattering, estimated as the odds ratio between ripe flowers that still had seeds attached versus flowers that did not have any seeds left at the time of harvest, was available for five cultivation lineages. We related the ratio of the number of flowers with seeds to the number of flowers without seeds as response variable to the generation as explanatory variable in a generalized linear model with quasibinomial error distribution. To illustrate the magnitudes of the changes in each response variable, we calculated effect sizes (i.e. changes per generation) as the proportions of the data range of the F0, calculated as the standard deviation times four.

## Acknowledgments

We are grateful to Saaten Zeller GmbH, Germany, and to Wilhelm Graiss, HBLFA Raumberg-Gumpenstein, Austria, for providing the seeds. Our work was supported by the DFG projects LA4038/2-1 to AB and DU404/14-1 to WD. We thank Theresa Klein-Raufhake and Hannah Kalthoff for technical assistance.

## SUPPLEMENTARY INFORMATION

**Table S1.** Results of statistical models testing for changes of mean and of coefficient of variation in biomass, height, number of flowers and start of flowering across cultivated generations (see Figure 2 in the main text). *P*-values *<* 0.05 are in bold.

| Parameter | Mode | Estimate | Std.Error | P | Conditional R<sup>2</sup> | NumDf | DenDf | F |
| --- | --- | --- | --- | --- | --- | --- | --- | --- |
| Biomass [g] | Mean | 0.07 | 0.021 | **0.001** | 0.007 | 1559 | 10.55 | 16.29 |
| Height [cm] | Mean | 0.052 | 0.021 | **0.015** | 0.004 | 1587 | 5.92 | 5.92 |
| Number of flowers | Mean | 0.064 | 0.027 | **0.019** | 0.006 | 918 | 5.55 | 5.55 |
| Start of flowering [d] | Mean | -0.049 | 0.027 | 0.069 | 0.003 | 948 | 3.31 | 3.31 |
| Biomass [g] | Variation | -0.015 | 0.007 | **0.029** | 0.392 | 72.89 | 4.93 | 4.93 |
| Height [cm] | Variation | 0.023 | 0.012 | 0.065 | 0.733 | 67.08 | 3.52 | 3.52 |
| Number of flowers | Variation | -0.044 | 0.017 | **0.010** | 0.412 | 50.57 | 7.12 | 7.12 |
| Start of flowering [d] | Variation | -0.007 | 0.003 | **0.010** | 0.531 | 40.12 | 7.40 | 7.40 |

**Table S2.**
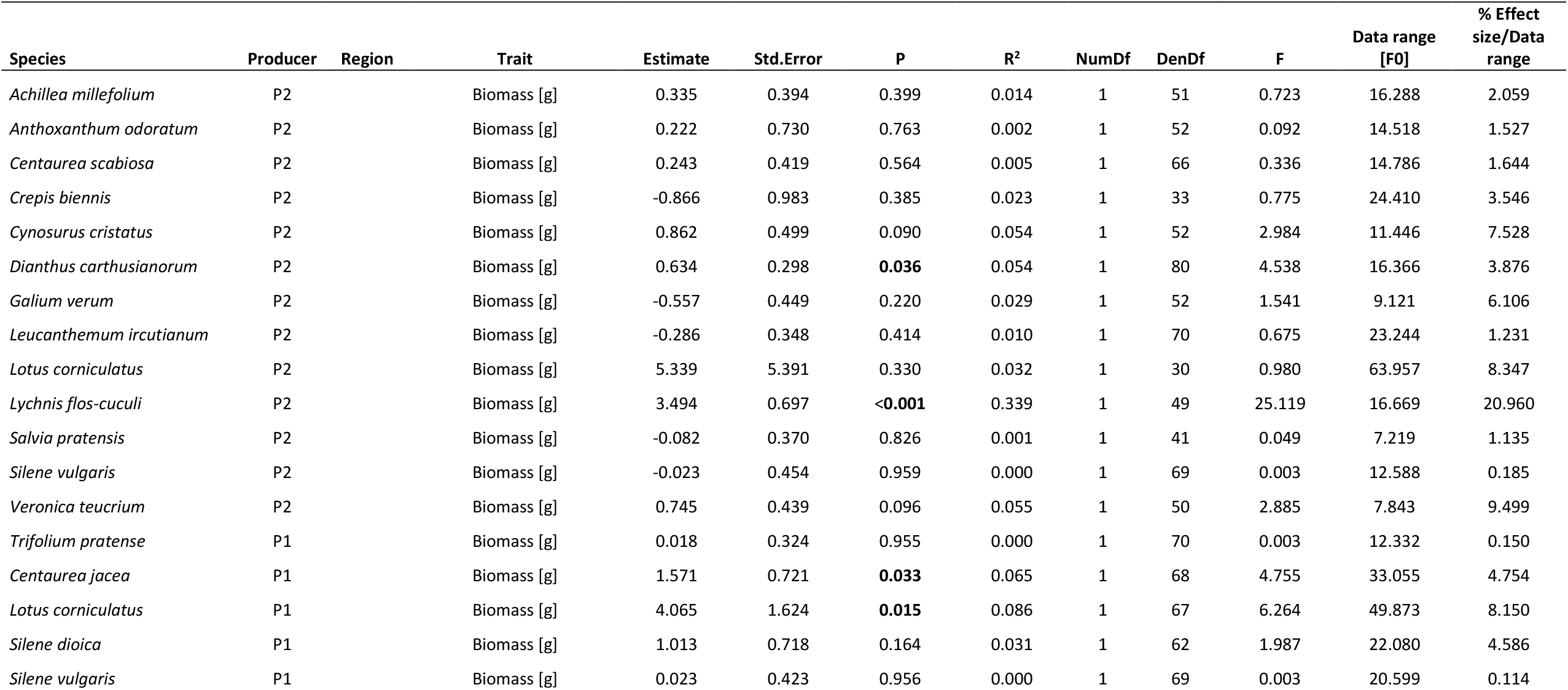
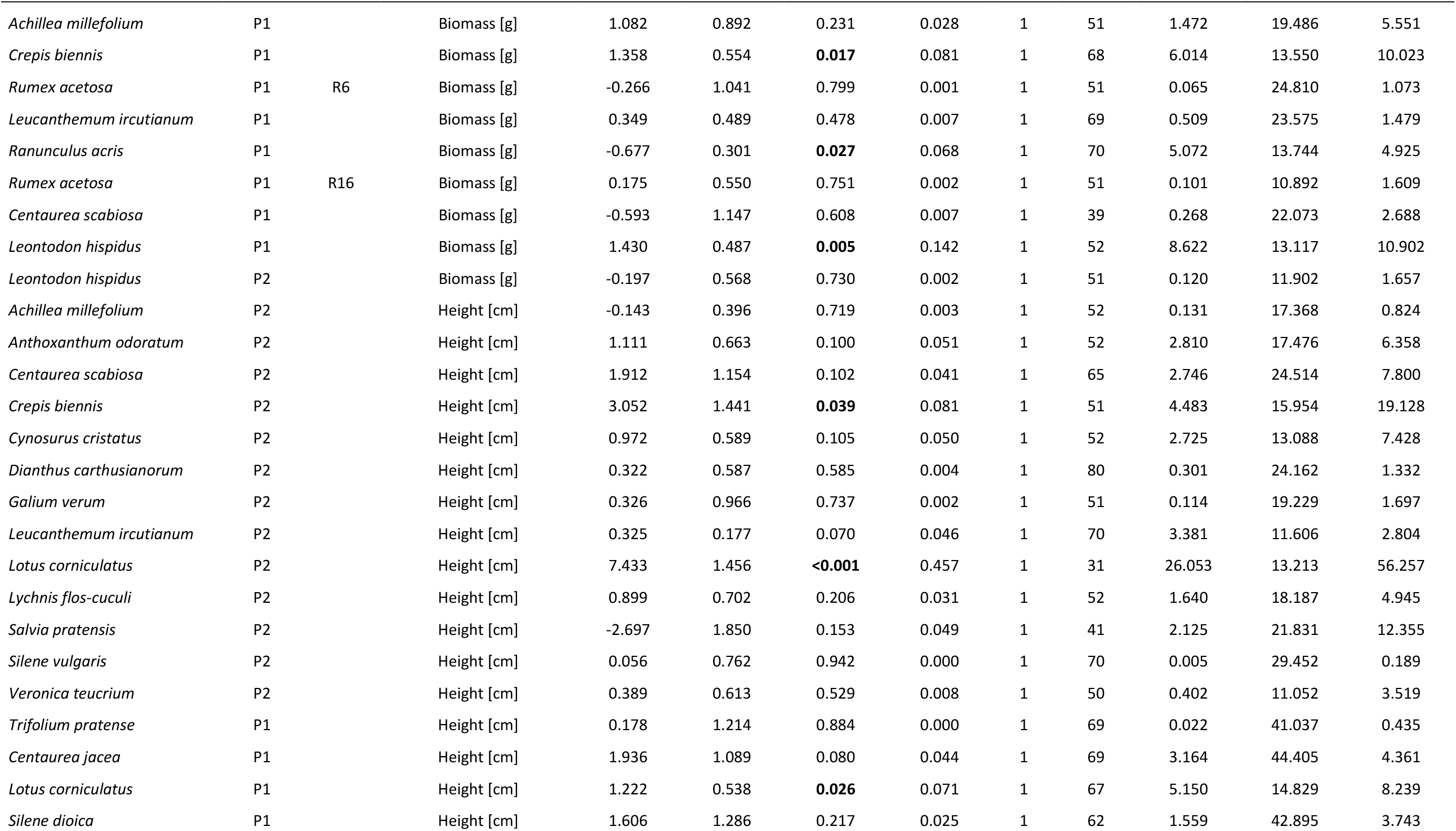
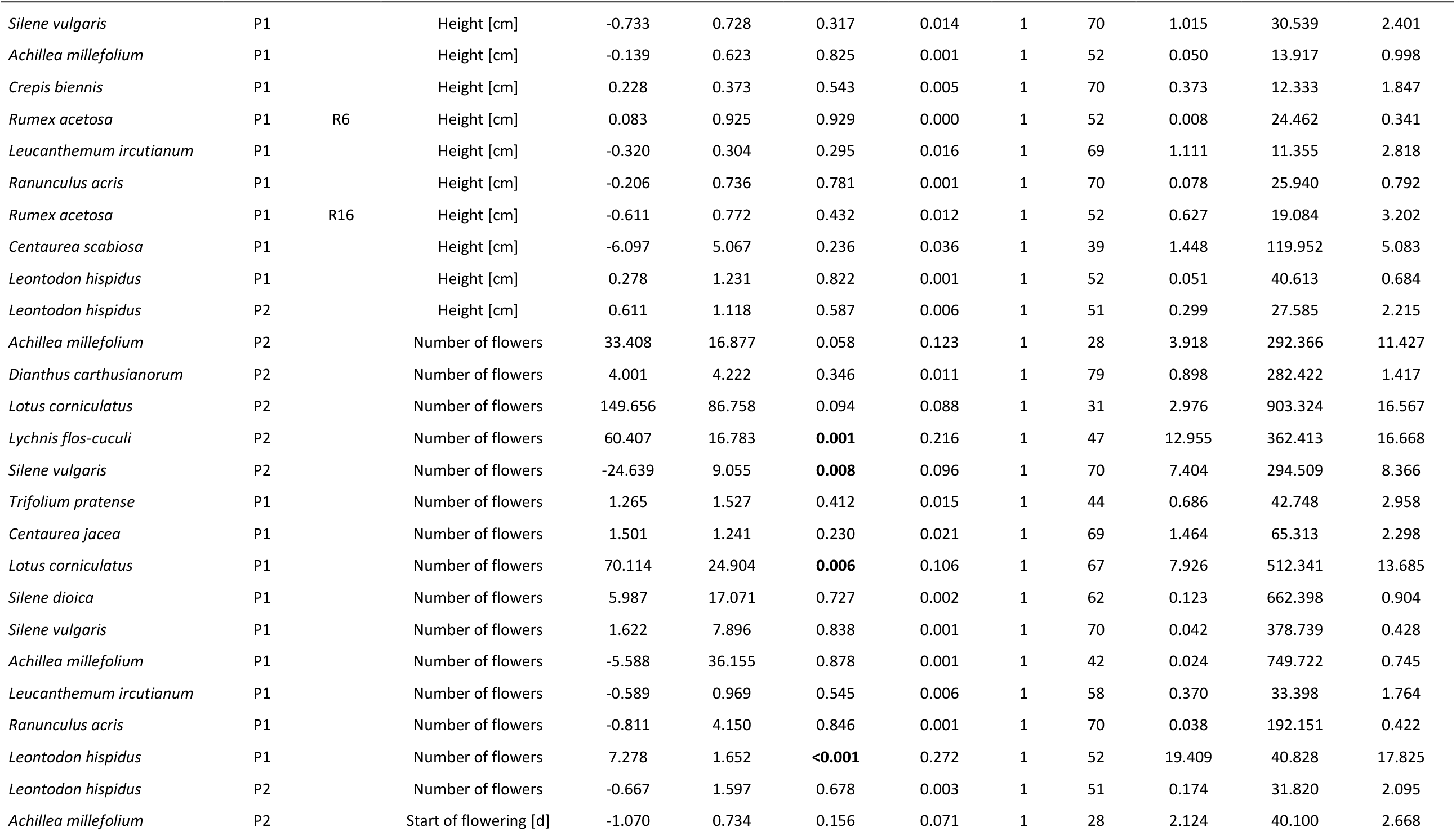
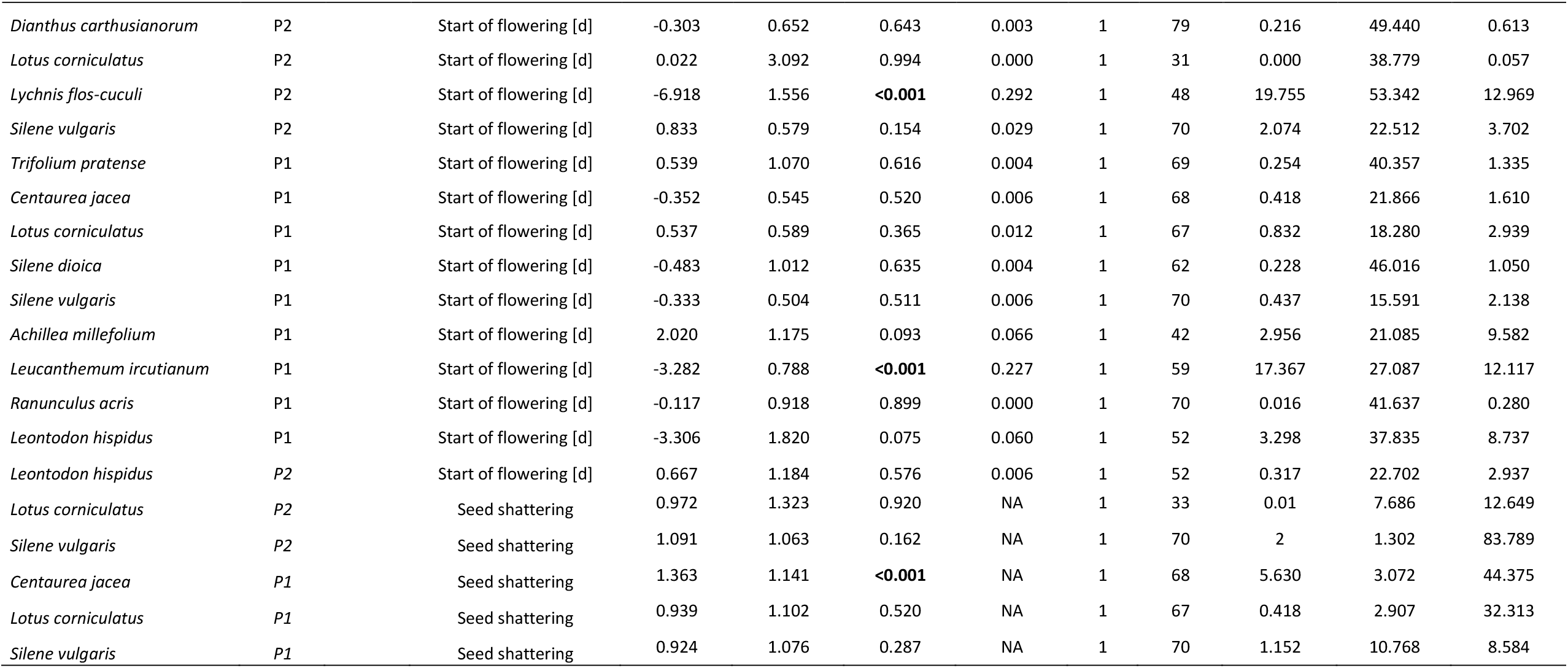
Results of models testing for changes in biomass, height, number of flowers and seed shattering within individual cultivation lineages. The “data range [F0]” was calculated as four-times standard deviation of the respective parameter in the F0. The last column shows the change per generation expressed as % of the data range among the plants from the wild collection (F0). Seed shattering is expressed as the odds of number of flowers that still had seeds at the end of harvest versus flowers that lost the seeds. *P*-values ≤ 0.05 are in bold.

| Species | Producer | Region | Trait | Estimate | Std.Error | P | R<sup>2</sup> | NumDf | DenDf | F | Data range [F0] | % Effect size/Data range |
| --- | --- | --- | --- | --- | --- | --- | --- | --- | --- | --- | --- | --- |
| *Achillea millefolium* | P2 |  | Biomass [g] | 0.335 | 0.394 | 0.399 | 0.014 | 1 | 51 | 0.723 | 16.288 | 2.059 |
| *Anthoxanthum odoratum* | P2 |  | Biomass [g] | 0.222 | 0.730 | 0.763 | 0.002 | 1 | 52 | 0.092 | 14.518 | 1.527 |
| *Centaurea scabiosa* | P2 |  | Biomass [g] | 0.243 | 0.419 | 0.564 | 0.005 | 1 | 66 | 0.336 | 14.786 | 1.644 |
| *Crepis biennis* | P2 |  | Biomass [g] | -0.866 | 0.983 | 0.385 | 0.023 | 1 | 33 | 0.775 | 24.410 | 3.546 |
| *Cynosurus cristatus* | P2 |  | Biomass [g] | 0.862 | 0.499 | 0.090 | 0.054 | 1 | 52 | 2.984 | 11.446 | 7.528 |
| *Dianthus carthusianorum* | P2 |  | Biomass [g] | 0.634 | 0.298 | **0.036** | 0.054 | 1 | 80 | 4.538 | 16.366 | 3.876 |
| *Galium verum* | P2 |  | Biomass [g] | -0.557 | 0.449 | 0.220 | 0.029 | 1 | 52 | 1.541 | 9.121 | 6.106 |
| *Leucanthemum ircutianum* | P2 |  | Biomass [g] | -0.286 | 0.348 | 0.414 | 0.010 | 1 | 70 | 0.675 | 23.244 | 1.231 |
| *Lotus corniculatus* | P2 |  | Biomass [g] | 5.339 | 5.391 | 0.330 | 0.032 | 1 | 30 | 0.980 | 63.957 | 8.347 |
| *Lychnis flos-cuculi* | P2 |  | Biomass [g] | 3.494 | 0.697 | **<0.001** | 0.339 | 1 | 49 | 25.119 | 16.669 | 20.960 |
| *Salvia pratensis* | P2 |  | Biomass [g] | -0.082 | 0.370 | 0.826 | 0.001 | 1 | 41 | 0.049 | 7.219 | 1.135 |
| *Silene vulgaris* | P2 |  | Biomass [g] | -0.023 | 0.454 | 0.959 | 0.000 | 1 | 69 | 0.003 | 12.588 | 0.185 |
| *Veronica teucrium* | P2 |  | Biomass [g] | 0.745 | 0.439 | 0.096 | 0.055 | 1 | 50 | 2.885 | 7.843 | 9.499 |
| *Trifolium pratense* | P1 |  | Biomass [g] | 0.018 | 0.324 | 0.955 | 0.000 | 1 | 70 | 0.003 | 12.332 | 0.150 |
| *Centaurea jacea* | P1 |  | Biomass [g] | 1.571 | 0.721 | **0.033** | 0.065 | 1 | 68 | 4.755 | 33.055 | 4.754 |
| *Lotus corniculatus* | P1 |  | Biomass [g] | 4.065 | 1.624 | **0.015** | 0.086 | 1 | 67 | 6.264 | 49.873 | 8.150 |
| *Silene dioica* | P1 |  | Biomass [g] | 1.013 | 0.718 | 0.164 | 0.031 | 1 | 62 | 1.987 | 22.080 | 4.586 |
| *Silene vulgaris* | P1 |  | Biomass [g] | 0.023 | 0.423 | 0.956 | 0.000 | 1 | 69 | 0.003 | 20.599 | 0.114 |
| *Achillea millefolium* | P1 |  | Biomass [g] | 1.082 | 0.892 | 0.231 | 0.028 | 1 | 51 | 1.472 | 19.486 | 5.551 |
| *Crepis biennis* | P1 |  | Biomass [g] | 1.358 | 0.554 | **0.017** | 0.081 | 1 | 68 | 6.014 | 13.550 | 10.023 |
| *Rumex acetosa* | P1 | R6 | Biomass [g] | -0.266 | 1.041 | 0.799 | 0.001 | 1 | 51 | 0.065 | 24.810 | 1.073 |
| *Leucanthemum ircutianum* | P1 |  | Biomass [g] | 0.349 | 0.489 | 0.478 | 0.007 | 1 | 69 | 0.509 | 23.575 | 1.479 |
| *Ranunculus acris* | P1 |  | Biomass [g] | -0.677 | 0.301 | **0.027** | 0.068 | 1 | 70 | 5.072 | 13.744 | 4.925 |
| *Rumex acetosa* | P1 | R16 | Biomass [g] | 0.175 | 0.550 | 0.751 | 0.002 | 1 | 51 | 0.101 | 10.892 | 1.609 |
| *Centaurea scabiosa* | P1 |  | Biomass [g] | -0.593 | 1.147 | 0.608 | 0.007 | 1 | 39 | 0.268 | 22.073 | 2.688 |
| *Leontodon hispidus* | P1 |  | Biomass [g] | 1.430 | 0.487 | **0.005** | 0.142 | 1 | 52 | 8.622 | 13.117 | 10.902 |
| *Leontodon hispidus* | P2 |  | Biomass [g] | -0.197 | 0.568 | 0.730 | 0.002 | 1 | 51 | 0.120 | 11.902 | 1.657 |
| *Achillea millefolium* | P2 |  | Height [cm] | -0.143 | 0.396 | 0.719 | 0.003 | 1 | 52 | 0.131 | 17.368 | 0.824 |
| *Anthoxanthum odoratum* | P2 |  | Height [cm] | 1.111 | 0.663 | 0.100 | 0.051 | 1 | 52 | 2.810 | 17.476 | 6.358 |
| *Centaurea scabiosa* | P2 |  | Height [cm] | 1.912 | 1.154 | 0.102 | 0.041 | 1 | 65 | 2.746 | 24.514 | 7.800 |
| *Crepis biennis* | P2 |  | Height [cm] | 3.052 | 1.441 | **0.039** | 0.081 | 1 | 51 | 4.483 | 15.954 | 19.128 |
| *Cynosurus cristatus* | P2 |  | Height [cm] | 0.972 | 0.589 | 0.105 | 0.050 | 1 | 52 | 2.725 | 13.088 | 7.428 |
| *Dianthus carthusianorum* | P2 |  | Height [cm] | 0.322 | 0.587 | 0.585 | 0.004 | 1 | 80 | 0.301 | 24.162 | 1.332 |
| *Galium verum* | P2 |  | Height [cm] | 0.326 | 0.966 | 0.737 | 0.002 | 1 | 51 | 0.114 | 19.229 | 1.697 |
| *Leucanthemum ircutianum* | P2 |  | Height [cm] | 0.325 | 0.177 | 0.070 | 0.046 | 1 | 70 | 3.381 | 11.606 | 2.804 |
| *Lotus corniculatus* | P2 |  | Height [cm] | 7.433 | 1.456 | **<0.001** | 0.457 | 1 | 31 | 26.053 | 13.213 | 56.257 |
| *Lychnis flos-cuculi* | P2 |  | Height [cm] | 0.899 | 0.702 | 0.206 | 0.031 | 1 | 52 | 1.640 | 18.187 | 4.945 |
| *Salvia pratensis* | P2 |  | Height [cm] | -2.697 | 1.850 | 0.153 | 0.049 | 1 | 41 | 2.125 | 21.831 | 12.355 |
| *Silene vulgaris* | P2 |  | Height [cm] | 0.056 | 0.762 | 0.942 | 0.000 | 1 | 70 | 0.005 | 29.452 | 0.189 |
| *Veronica teucrium* | P2 |  | Height [cm] | 0.389 | 0.613 | 0.529 | 0.008 | 1 | 50 | 0.402 | 11.052 | 3.519 |
| *Trifolium pratense* | P1 |  | Height [cm] | 0.178 | 1.214 | 0.884 | 0.000 | 1 | 69 | 0.022 | 41.037 | 0.435 |
| *Centaurea jacea* | P1 |  | Height [cm] | 1.936 | 1.089 | 0.080 | 0.044 | 1 | 69 | 3.164 | 44.405 | 4.361 |
| *Lotus corniculatus* | P1 |  | Height [cm] | 1.222 | 0.538 | **0.026** | 0.071 | 1 | 67 | 5.150 | 14.829 | 8.239 |
| *Silene dioica* | P1 |  | Height [cm] | 1.606 | 1.286 | 0.217 | 0.025 | 1 | 62 | 1.559 | 42.895 | 3.743 |
| *Silene vulgaris* | P1 |  | Height [cm] | -0.733 | 0.728 | 0.317 | 0.014 | 1 | 70 | 1.015 | 30.539 | 2.401 |
| *Achillea millefolium* | P1 |  | Height [cm] | -0.139 | 0.623 | 0.825 | 0.001 | 1 | 52 | 0.050 | 13.917 | 0.998 |
| *Crepis biennis* | P1 |  | Height [cm] | 0.228 | 0.373 | 0.543 | 0.005 | 1 | 70 | 0.373 | 12.333 | 1.847 |
| *Rumex acetosa* | P1 | R6 | Height [cm] | 0.083 | 0.925 | 0.929 | 0.000 | 1 | 52 | 0.008 | 24.462 | 0.341 |
| *Leucanthemum ircutianum* | P1 |  | Height [cm] | -0.320 | 0.304 | 0.295 | 0.016 | 1 | 69 | 1.111 | 11.355 | 2.818 |
| *Ranunculus acris* | P1 |  | Height [cm] | -0.206 | 0.736 | 0.781 | 0.001 | 1 | 70 | 0.078 | 25.940 | 0.792 |
| *Rumex acetosa* | P1 | R16 | Height [cm] | -0.611 | 0.772 | 0.432 | 0.012 | 1 | 52 | 0.627 | 19.084 | 3.202 |
| *Centaurea scabiosa* | P1 |  | Height [cm] | -6.097 | 5.067 | 0.236 | 0.036 | 1 | 39 | 1.448 | 119.952 | 5.083 |
| *Leontodon hispidus* | P1 |  | Height [cm] | 0.278 | 1.231 | 0.822 | 0.001 | 1 | 52 | 0.051 | 40.613 | 0.684 |
| *Leontodon hispidus* | P2 |  | Height [cm] | 0.611 | 1.118 | 0.587 | 0.006 | 1 | 51 | 0.299 | 27.585 | 2.215 |
| *Achillea millefolium* | P2 |  | Number of flowers | 33.408 | 16.877 | 0.058 | 0.123 | 1 | 28 | 3.918 | 292.366 | 11.427 |
| *Dianthus carthusianorum* | P2 |  | Number of flowers | 4.001 | 4.222 | 0.346 | 0.011 | 1 | 79 | 0.898 | 282.422 | 1.417 |
| *Lotus corniculatus* | P2 |  | Number of flowers | 149.656 | 86.758 | 0.094 | 0.088 | 1 | 31 | 2.976 | 903.324 | 16.567 |
| *Lychnis flos-cuculi* | P2 |  | Number of flowers | 60.407 | 16.783 | **0.001** | 0.216 | 1 | 47 | 12.955 | 362.413 | 16.668 |
| *Silene vulgaris* | P2 |  | Number of flowers | -24.639 | 9.055 | **0.008** | 0.096 | 1 | 70 | 7.404 | 294.509 | 8.366 |
| *Trifolium pratense* | P1 |  | Number of flowers | 1.265 | 1.527 | 0.412 | 0.015 | 1 | 44 | 0.686 | 42.748 | 2.958 |
| *Centaurea jacea* | P1 |  | Number of flowers | 1.501 | 1.241 | 0.230 | 0.021 | 1 | 69 | 1.464 | 65.313 | 2.298 |
| *Lotus corniculatus* | P1 |  | Number of flowers | 70.114 | 24.904 | **0.006** | 0.106 | 1 | 67 | 7.926 | 512.341 | 13.685 |
| *Silene dioica* | P1 |  | Number of flowers | 5.987 | 17.071 | 0.727 | 0.002 | 1 | 62 | 0.123 | 662.398 | 0.904 |
| *Silene vulgaris* | P1 |  | Number of flowers | 1.622 | 7.896 | 0.838 | 0.001 | 1 | 70 | 0.042 | 378.739 | 0.428 |
| *Achillea millefolium* | P1 |  | Number of flowers | -5.588 | 36.155 | 0.878 | 0.001 | 1 | 42 | 0.024 | 749.722 | 0.745 |
| *Leucanthemum ircutianum* | P1 |  | Number of flowers | -0.589 | 0.969 | 0.545 | 0.006 | 1 | 58 | 0.370 | 33.398 | 1.764 |
| *Ranunculus acris* | P1 |  | Number of flowers | -0.811 | 4.150 | 0.846 | 0.001 | 1 | 70 | 0.038 | 192.151 | 0.422 |
| *Leontodon hispidus* | P1 |  | Number of flowers | 7.278 | 1.652 | **<0.001** | 0.272 | 1 | 52 | 19.409 | 40.828 | 17.825 |
| *Leontodon hispidus* | P2 |  | Number of flowers | -0.667 | 1.597 | 0.678 | 0.003 | 1 | 51 | 0.174 | 31.820 | 2.095 |
| *Achillea millefolium* | P2 |  | Start of flowering [d] | -1.070 | 0.734 | 0.156 | 0.071 | 1 | 28 | 2.124 | 40.100 | 2.668 |
| *Dianthus carthusianorum* | P2 |  | Start of flowering [d] | -0.303 | 0.652 | 0.643 | 0.003 | 1 | 79 | 0.216 | 49.440 | 0.613 |
| *Lotus corniculatus* | P2 |  | Start of flowering [d] | 0.022 | 3.092 | 0.994 | 0.000 | 1 | 31 | 0.000 | 38.779 | 0.057 |
| *Lychnis flos-cuculi* | P2 |  | Start of flowering [d] | -6.918 | 1.556 | **<0.001** | 0.292 | 1 | 48 | 19.755 | 53.342 | 12.969 |
| *Silene vulgaris* | P2 |  | Start of flowering [d] | 0.833 | 0.579 | 0.154 | 0.029 | 1 | 70 | 2.074 | 22.512 | 3.702 |
| *Trifolium pratense* | P1 |  | Start of flowering [d] | 0.539 | 1.070 | 0.616 | 0.004 | 1 | 69 | 0.254 | 40.357 | 1.335 |
| *Centaurea jacea* | P1 |  | Start of flowering [d] | -0.352 | 0.545 | 0.520 | 0.006 | 1 | 68 | 0.418 | 21.866 | 1.610 |
| *Lotus corniculatus* | P1 |  | Start of flowering [d] | 0.537 | 0.589 | 0.365 | 0.012 | 1 | 67 | 0.832 | 18.280 | 2.939 |
| *Silene dioica* | P1 |  | Start of flowering [d] | -0.483 | 1.012 | 0.635 | 0.004 | 1 | 62 | 0.228 | 46.016 | 1.050 |
| *Silene vulgaris* | P1 |  | Start of flowering [d] | -0.333 | 0.504 | 0.511 | 0.006 | 1 | 70 | 0.437 | 15.591 | 2.138 |
| *Achillea millefolium* | P1 |  | Start of flowering [d] | 2.020 | 1.175 | 0.093 | 0.066 | 1 | 42 | 2.956 | 21.085 | 9.582 |
| *Leucanthemum ircutianum* | P1 |  | Start of flowering [d] | -3.282 | 0.788 | **<0.001** | 0.227 | 1 | 59 | 17.367 | 27.087 | 12.117 |
| *Ranunculus acris* | P1 |  | Start of flowering [d] | -0.117 | 0.918 | 0.899 | 0.000 | 1 | 70 | 0.016 | 41.637 | 0.280 |
| *Leontodon hispidus* | P1 |  | Start of flowering [d] | -3.306 | 1.820 | 0.075 | 0.060 | 1 | 52 | 3.298 | 37.835 | 8.737 |
| *Leontodon hispidus* | P2 |  | Start of flowering [d] | 0.667 | 1.184 | 0.576 | 0.006 | 1 | 52 | 0.317 | 22.702 | 2.937 |
| *Lotus corniculatus* | P2 |  | Seed shattering | 0.972 | 1.323 | 0.920 | NA | 1 | 33 | 0.01 | 7.686 | 12.649 |
| *Silene vulgaris* | P2 |  | Seed shattering | 1.091 | 1.063 | 0.162 | NA | 1 | 70 | 2 | 1.302 | 83.789 |
| *Centaurea jacea* | P1 |  | Seed shattering | 1.363 | 1.141 | **<0.001** | NA | 1 | 68 | 5.630 | 3.072 | 44.375 |
| *Lotus corniculatus* | P1 |  | Seed shattering | 0.939 | 1.102 | 0.520 | NA | 1 | 67 | 0.418 | 2.907 | 32.313 |
| *Silene vulgaris* | P1 |  | Seed shattering | 0.924 | 1.076 | 0.287 | NA | 1 | 70 | 1.152 | 10.768 | 8.584 |

**Figure S1.**
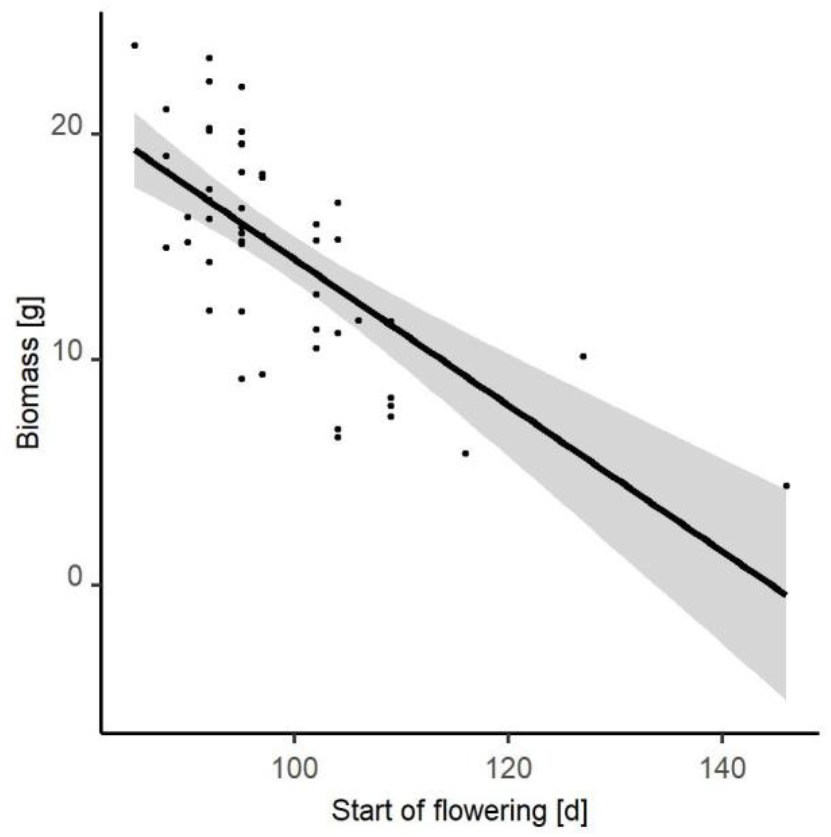
The relationship between flowering time and biomass in *Lychnis floss-cuculi*, as estimated from a linear model. The relationship is significant at *P* < 0.01.

